# Redox regulation of 2-carboxy-D-arabinitol 1-phosphate phosphatase in Fabaceae is associated with high Rubisco dark inhibition

**DOI:** 10.64898/2026.11.05.764501

**Authors:** Connor Nehls-Ramos, Elizabete Carmo-Silva, Douglas J. Orr

**Author notes:** **Supplementary Data:** In file or otherwise accessible by Github: cwnehls/NehlsRamos_etal_JxB_2027: Supplementary data and code for the publication Nehls-Ramos *et al*. 2027.

## Abstract

Improving Rubisco regulation is an avenue for improving crop output. Minimizing the buildup of Rubisco inhibitors can allow improved catalytic activity and CO_2_ assimilation. The enzyme 2-carboxy-D-arabinitol 1-phosphate phosphatase (CA1Pase) degrades certain Rubisco inhibitors, yet the mechanisms for its regulation, including the function of its C domain, are not well understood. In this study we analysed *Vigna unguiculata* (cowpea) CA1Pase *in silico* and *in vitro* for structural regions and motifs related to regulation of phosphatase activity. We identified a highly conserved cysteine pair motif tied to redox regulation and potential homodimer formation. Variations in one or both cysteine residues in this motif across species correlate with lower dark inhibitor accumulation levels as compared to related species. The C domain is identified as having a regulatory function and potential evolutionary origins are explored. These insights may inform future efforts toward optimising Rubisco’s complex regulatory processes including inhibitor degradation.

**Highlight:** A cysteine pair motif confers redox regulation to a chloroplast phosphatase that degrades Rubisco inhibitors. *In silico* and *in vitro* analyses suggest this mechanism correlates with Rubisco dark inhibition in Fabaceae.

## Introduction

One strategy to increase crop output is by improving photosynthesis through fixing inefficiencies associated with Rubisco, the imperfect enzyme essential to carbon fixation in photosynthesis (Parry *et al*., 2013; Croce *et al*., 2024). The activity of Rubisco is regulated in response to environmental factors such as light availability, which is impacted within the crop canopy by cloud cover, wind speed and sunflecks frequency and duration, as well as sun movement across the day (Yamori, 2016; Slattery *et al*., 2018). Despite advances in understanding the complex regulation of Rubisco activity, there remain many knowledge gaps (Parry *et al*., 2008; Leister *et al*., 2023; Orr *et al*., 2023; Amaral *et al*., 2024; Pasch *et al*., 2025). While it has long been known that certain sugar phosphate inhibitors bind tightly to Rubisco catalytic sites and prevent catalysis throughout the day, it is less clear what factors regulate their abundance and degradation (Parry *et al*., 2008; Orr *et al*., 2023; Pasch *et al*., 2025).

For many but not all crops, accumulation of nighttime Rubisco inhibitors, thought to be primarily 2-carboxy-D-arabinitol 1-phosphate (CA1P), has been reported when leaves experience an extended period of low light or darkness, as seen in a recent compilation of dark inhibition across angiosperms (Nehls-Ramos *et al*., 2026). For cowpea specifically, this dark inhibitor has been found to accumulate to significant amounts (Servaites *et al*., 1986). There are two known chloroplast-localised phosphatases which dephosphorylate Rubisco inhibitors, 2-carboxy-D-arabinitol 1-phosphate phosphatase (CA1Pase; EC: 3.1.3.63) and xylulose-1,5-bisphosphate phosphatase (Parry *et al*., 2008; Bracher *et al*., 2015). CA1Pase dephosphorylates not only the diurnal inhibitor CA1P, but also Rubisco misfire products including D-glycero-2,3-pentodiulose-1,5-bisphosphate (PDBP) after release by Rubisco activase (Parry *et al*., 2008; Andralojc *et al*., 2012).

CA1Pase is a histidine phosphatase and part of the ancient phosphoglycerate mutase (PGM) family. This enzyme is believed to reside solely in the chloroplast stroma (Salvucci *et al*., 1988; Holbrook *et al*., 1989; Gutteridge & Julien, 1989; Moore *et al*., 1995; Bayer *et al*., 2011). The mature enzyme in flowering plants typically ranges between 440-460 amino acids in length, with a monomeric size near 50kDa. A previous study identified two distinct domains of similar length in wheat and Arabidopsis (Andralojc *et al*., 2012). The N domain contains the histidine phosphatase catalytic region, in which reside five main catalytic residues: two arginines (R), two histidines (H), and a glutamic acid (E) (Rigden, 2007; Andralojc *et al*., 2012). The role of the C domain has not been defined, though it has been previously hypothesised to be a phosphofructokinase-like domain (Andralojc *et al*., 2012).

Despite nearly 40 years of investigation, insights on the *in vivo* role and regulation of CA1P and CA1Pase are limited, especially regarding the intersection of enzyme activity regulation and structural biology (Salvucci *et al*., 1988; Holbrook *et al*., 1989; Gutteridge & Julien, 1989; Salvucci *et al*., 1989; Holbrook *et al*., 1991; Kingston Smith *et al*., 1992; Moore *et al*., 1995; Charlet *et al*., 1997; Heo & Holbrook 1999; Andralojc *et al*., 2012; Lobo *et al*., 2019). Regulation of CA1Pase has been directly investigated in five species: tobacco (*Nicotiana tabacum* & *N. rustica*), French bean (*Phaseolus vulgaris*), spinach (*Spinacia oleracea*), and wheat (*Triticum aestivum*), with some other investigations into the kinetics in other species (Table 1; Charlet *et al*., 1997). While there are large variations in dark inhibitor accumulation in plants, CA1Pase appears to be ubiquitous across angiosperms, and may be present in more ancient lineages including algae (Andralojc *et al*., 2012; Nehls-Ramos *et al*., 2026). An ancient CA1Pase would imply a role deeply entrenched in chloroplast and Rubisco homeostasis, which may explain why modifications to its abundance have caused unexpected shifts in Rubisco abundance in plants (Lobo *et al*., 2019). To better understand the importance of CA1Pase in the chloroplast as well as retained features in structure, its potential ancestral origins should be explored further.

**Table 1.**
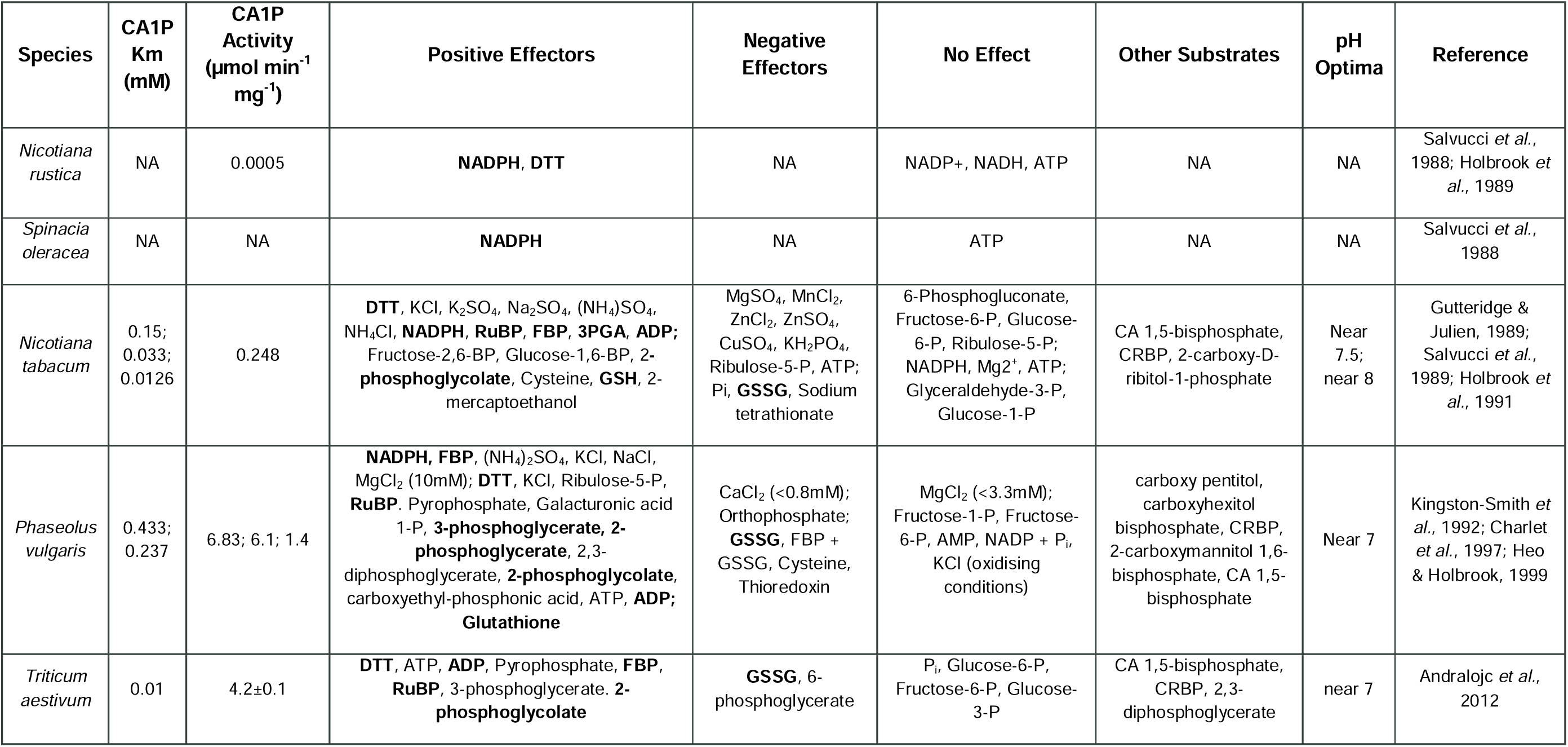
Available data characterising CA1Pase: Michaelis Menten constant (Km) for CA1P degradation; molecules that increase, crease or do not significantly affect CA1Pase activity; substrates in addition to CA1P; and pH optima. Bold indicates positive ectors identified in multiple studies or species. NA, not available.

| **Species** | **CA1P Km (mM)** | **CA1P Activity (µmol min<sup>-1</sup> mg<sup>-1</sup>)** | **Positive Effectors** | **Negative Effectors** | **No Effect** | **Other Substrates** | **pH Optima** | **Reference** |
|---|---|---|---|---|---|---|---|---|
| *Nicotiana rustica* | NA | 0.0005 | **NADPH**, **DTT** | NA | NADP+, NADH, ATP | NA | NA | Salvucci *et al.*, 1988; Holbrook *et al.*, 1989 |
| *Spinacia oleracea* | NA | NA | **NADPH** | NA | ATP | NA | NA | Salvucci *et al.*, 1988 |
| *Nicotiana tabacum* | 0.15; 0.033; 0.0126 | 0.248 | **DTT**, KCl, K<sub>2</sub>SO<sub>4</sub>, Na<sub>2</sub>SO<sub>4</sub>, (NH<sub>4</sub>)SO<sub>4</sub>, NH<sub>4</sub>Cl, **NADPH**, **RuBP**, **FBP**, **3PGA**, **ADP;** Fructose-2,6-BP, Glucose-1,6-BP, **2-phosphoglycolate**, Cysteine, **GSH**, 2-mercaptoethanol | MgSO<sub>4</sub>, MnCl<sub>2</sub>, ZnCl<sub>2</sub>, ZnSO<sub>4</sub>, CuSO<sub>4</sub>, KH<sub>2</sub>PO<sub>4</sub>, Ribulose-5-P, ATP; Pi, **GSSG**, Sodium tetrathionate | 6-Phosphogluconate, Fructose-6-P, Glucose-6-P, Ribulose-5-P; NADPH, Mg2<sup>+</sup>, ATP; Glyceraldehyde-3-P, Glucose-1-P | CA 1,5-bisphosphate, CRBP, 2-carboxy-D-ribitol-1-phosphate | Near 7.5; near 8 | Gutteridge & Julien, 1989; Salvucci *et al.*, 1989; Holbrook *et al.*, 1991 |
| *Phaseolus vulgaris* | 0.433; 0.237 | 6.83; 6.1; 1.4 | **NADPH, FBP**, (NH<sub>4</sub>)<sub>2</sub>SO<sub>4</sub>, KCl, NaCl, MgCl<sub>2</sub> (10mM); **DTT**, KCl, Ribulose-5-P, **RuBP**. Pyrophosphate, Galacturonic acid 1-P, **3-phosphoglycerate, 2-phosphoglycerate**, 2,3-diphosphoglycerate, **2-phosphoglycolate**, carboxyethyl-phosphonic acid, ATP, **ADP; Glutathione** | CaCl<sub>2</sub> (<0.8mM); Orthophosphate; **GSSG**, FBP + GSSG, Cysteine, Thioredoxin | MgCl<sub>2</sub> (<3.3mM); Fructose-1-P, Fructose-6-P, AMP, NADP + P<sub>i</sub>, KCl (oxidising conditions) | carboxy pentitol, carboxyhexitol bisphosphate, CRBP, 2-carboxymannitol 1,6-bisphosphate, CA 1,5-bisphosphate | Near 7 | Kingston-Smith *et al.*, 1992; Charlet *et al.*, 1997; Heo & Holbrook, 1999 |
| *Triticum aestivum* | 0.01 | 4.2±0.1 | **DTT**, ATP, **ADP**, Pyrophosphate, **FBP**, **RuBP**, 3-phosphoglycerate. **2-phosphoglycolate** | **GSSG**, 6-phosphoglycerate | P<sub>i</sub>, Glucose-6-P, Fructose-6-P, Glucose-3-P | CA 1,5-bisphosphate, CRBP, 2,3-diphosphoglycerate | near 7 | Andralojc *et al.*, 2012 |

While studies show conflicting results within and across species for factors effecting *in vitro* regulation of CA1Pase activity, some common trends are present (Table 1). CA1Pase shows a pH optima around or below 7.5, suggesting peak activity when chloroplasts are transitioning between darkness and high light (Trinh & Masuda, 2022). Several Calvin Benson Basham (CBB) cycle related metabolites have been shown to act as positive effectors of CA1Pase, consistent with a stimulation of activity as photosynthetic metabolism increases in the light (Table 1). One such effector is fructose-1,6-bisphosphate (FBP), a known precursor for the formation of 2-carboxy-D-arabinitol (CA) and CA1P (Andralojc *et al*., 2002). Evidence suggests CA1Pase partially deactivates in oxidising conditions and reactivates in reducing conditions (Salvucci *et al*., 1988; Holbrook *et al*., 1989; Gutteridge & Julien, 1989; Salvucci *et al*., 1989; Holbrook *et al*., 1991; Kingston Smith *et al*., 1992; Charlet *et al*., 1997; Heo & Holbrook, 1999). This has been hypothesised to occur due to a key pair of cysteine residues, which remain unidentified despite availability of CA1Pase sequences more than a decade later (Heo & Holbrook,1999; Andralojc *et al*., 2012).

In the chloroplast, redox regulation is key to linking photosynthetic metabolism with electron transport chain activity during photosynthesis (Yu *et al*., 2020; Yoshida & Hisabori, 2023; Müller-Schüssele, 2024). Several CBB enzymes, including Rubisco activase (Rca) and fructose-1,6-bisphophatase, have been shown to be subject to this type of regulation (Portis *et al*., 2008; Gütle *et al*., 2016). During active photosynthesis, the electron transport chain transfers photon energy into chemical energy by generating ATP and a proton gradient via photosystem II, as well as allowing for the generation of NADPH from NADP+ in photosystem I (Yoshida & Hisabori, 2023; Trinh & Masuda, 2022). The NADPH generated allows for the reduction of thioredoxin (Trx) and glutathione (GSH)—or homoglutathione (hGSH) in Fabaceae—which in turn regulates redox sensitive enzymes through the breaking of key disulphide bonds and the formation of mixed disulphides by S-glutathionylation (Musaogullari & Chai, 2020). These stromal reducing agents also buffer for reactive oxygen species (ROS), such as H_2_O_2_, which are generated during photosynthesis and via cell signalling pathways (Foyer & Hanke, 2022). During dark periods, the reducing potential of the stroma greatly diminishes, resulting in the reversible oxidation of stromal enzyme cysteine residues (Müller-Schüssele *et al*., 2020; Cejudo *et al*., 2021). This cycling of oxidising (dark and low light) and reducing (light) conditions allows for a cyclical regulation of photosynthetic enzymes.

In this study we elucidate the mechanism for redox regulation in CA1Pase and clarify the function of the regulatory C domain, using CA1Pase from the staple legume cowpea (*Vigna unguiculata*). We use a combination of *in silico* sequence, predicted structure analyses, *in vitro* activity and oligomerisation measurements of recombinant cowpea CA1Pase (VuCA1Pase). Key cysteines were identified *in silico* and mutated to investigate their effect on redox regulation of CA1Pase activity. We propose that oxidation of this cysteine pair motif (CXXXXC) in CA1Pase is required for the high levels of Rubisco dark inhibition observed in flowering plants.

## Materials and methods

### CA1Pase sequences & identification

The CA1Pase sequence was sourced from the cowpea genome (Phytozome 13 database; Goodstein *et al*., 2012) for *Vigna unguiculata* 1.2 (ViGun11g130800.1.10) using known CA1Pase sequences from *Arabidopsis thaliana* and *Triticum aestivum* (Andralojc *et al*., 2012). To generate the mature sequence, the chloroplast transit peptide (cTP) predicted by Andralojc *et al*. (2012) was removed preceding the “KRVV…” motif. Key residues are numbered throughout using the mature protein sequence of cowpea CA1Pase (VuCA1Pase). Homologous CA1Pase sequences were further identified by performing a NCBI BLAST search against ViGun11g130800.1.10 to gather the top 1000 sequences related to the cowpea CA1Pase sequence, including searches for sequences identified in Uniprot with “2-carboxy-D-arabinitol 1-phosphatase” as the protein name (Sayers *et al*., 2025; The UniProt Consortium, 2025). The resulting sequences were curated for length between 400-600 amino acids, aligned using Geneious MUSCLE alignment and manually trimmed to remove those lacking the RHGQSTWN catalytic motif or the R/CXG and XXNXTXHL motifs unique to the Regulatory domain—highly conserved across all complete sequences. Using a consensus tree alignment, the sequences grouped into three main branches: cyanobacteria and chlorophytes, bryophytes, and all other streptophytes. The latter contains a branch with all angiosperm sequences (Supplementary Dataset S1).

### Predicted structure modelling and visualisation

Structural predictions were performed using Deepmind AlphaFold 3, with comparisons against D-I-Tasser modelling (Abramson *et al*., 2024; Zheng *et al*., 2026). Multimeric structures, including ligand interactions, were also modelled in AlphaFold 3. Similar predicted structures for the monomer and dimer forms of VuCA1Pase can be found with cTP included on the AlphaFold DB for *Vigna unguiculata* (AF-A0A4D6LHI6-F1-v6; AF-0000000079567473-v1; Bertoni *et al*., 2026). Individual predicted structures depicting model confidence, including predicted local distance difference test, can be found in Supplementary Fig. S1 and Supplementary Datasets S2 and S3. Structures were visualised and annotated in PyMol (version 2.5.7, Schrödinger, LLC., USA; www.pymol.org). Sequence alignments and trees were generated using Geneious (Geneious 9.1.8; www.geneious.com).

### Cloning of CA1Pase variants and expression vector creation

The mature VuCA1Pase sequence was inserted into a 6xHis TEV vector (pETHisTEVGG) (Gjindali *et al*., 2025). The domain boundary for designing the regulatory domain only variant, VuC_207-445_, was determined using D-I-Tasser-modelling (Supplementary Table S4). Modified cysteine variants are based on the 6xHis TEV VuCA1Pase sequence with one or both cysteine residues at C292 and C297 changed to Alanine.

To create the expression construct for the wild type form of Cowpea CA1Pase (Vigun11g130800.1), the oligonucleotide sequence, as well as the regulatory domain variant, was ordered from Aruru Molecular (United Kingdom) and amplified using primers ordered from Integrated DNA Technologies. The amplified sequence was inserted into the pETHisTEVGG golden gate compatible plasmid and expressed in BL21(DE3)pLysS *E. coli* with expression inducible T7 promoter. The Cysteine mutant variants were generated using the QuikChange II Site-Directed Mutagenesis Kit (200523; Agilent Technologies, Inc., Santa Clara, California, US; www.agilent.com), as per manufacturer specifications. Sequence inserted expression vectors were verified by Plasmidsaurus Whole Plasmid Sequencing (Oxford Nanopore, R10.4.1; Plasmidsaurus, UK; www.plasmidsaurus.com) and Source Bioscience (Nanopore Whole Plasmid Sequencing; Cambridge, UK; www.sourcebioscience.com).

### CA1Pase expression, purification and quantification

Transformed BL21(DE3)pLysS cells were expressed and the recombinant protein purified using the same method as per Gjindali *et al*. (2025). Purified protein was stored in 50 mM HEPES, pH 8.0, and 5% glycerol and aliquoted prior to flash freezing. Total soluble protein (TSP) was determined using the Bradford assay as well as 280 nm absorption via a CLARIOstar Plus spectrophotometer with an LVis plate (BMG Labtech, Ortenberg, Germany; www.bmglabtech.com) (Perdomo *et al*., 2018; Supplementary Dataset S5). For soluble protein calculations determined from the absorbance at 280 nm absorption, absorption coefficients for the 6xHis TEV tagged proteins were 63940 M^-1^ cm^-1^ for VuCA1Pase, 33460 M^-1^ cm^-1^ for VuC_1-206_, and 31970 M^-1^ cm^-1^ for VuC_207-445_ as determined using the Expasy Prot Param webtool (Gasteiger *et al*., 2005, https://web.expasy.org/protparam/). Protein purity determination by SDS-PAGE gel and western blot were performed as per Perdomo *et al*. (2018). Western blots used a mouse anti His antibody (Bio-Rad Cat#MCA1396D800GA, RRID: AB_11152596; www.bio-rad-antibodies.com. All gels were imaged using a Licor Odyssey Fc Imager (Li-Cor Bioscience, USA; www.licorbio.com). Purity of protein was quantified by densitometry using the Licor software ImageStudio comparing the peak density of the expressed protein in the Coomassie stained SDS-Page gels as a percentage of the remaining 700 nm absorption in the lane. Purity for all assayed proteins was >50% (Supplementary Fig. S2; Supplementary Table S2).

### CA1Pase activity assays

All reagents in this section and below were sourced from Sigma-Aldrich (UK; www.sigmaaldrich.com) unless otherwise specified. Activity was quantified using a modified version of a malachite phosphate detection assay (Martínez Gache *et al*., 2020). The colour detection reagent used (Supplementary Fig. S3) was modified to use malachite chloride in place of malachite green carbinol chloride (the latter being unavailable from commercial sources at the time). Final concentrations of reagents were: 14.24 mM ammonium molybdate tetrahydrate (A7302), 501.5 µM malachite chloride (38800), and 1.19 mM pluronic F68 in a solution of 16.8% (v/v) HClO_4_ (1.00517.0250), 0.5% (v/v) absolute ethanol (Fisher-Scientific, USA, E/0650DF/17), and the remainder milli-Q water. The solution was then filtered with ashless filter paper (Whatman, UK; www.cytivalifesciences.com) and stored in a brown polyethylene bottle. This solution is dark yellow-orange, clear, and can be used for up to one year.

Thawed protein was re-checked for TSP by A280, diluted in buffer (50 mM HEPES [Melford, H75030, UK], pH 8, and 200 mM NaCl [31434]), factoring for the purity of the sample, to a concentration of 1.186 μM of recombinant CA1Pase. A reaction buffer was prepared from a 10x stock (500 mM HEPES, pH 8.0, 50% glycerol w/v [G7757]) diluted in Mili-Q water. For assays involving glutathione, 200mM reduced (GSH) (G4251) and oxidised (GSSG) (G4376) glutathione stocks prepared in Mili-Q water were combined in differing ratios and included in the final buffer at a final concentration of 10mM. The sample buffer was added to a 96 well plate at 25°C on a plate incubator at a volume of 57 μL. In the same plate, a set of standards that contained 0.1-5 nmoles of pH 7.0 NaPi (0.4167 mM NaH_2_PO_4_ [S5011] and 0.5833 mM Na_2_HPO_4_ [S0876] for a 1 mM solution) were prepared and used to quantify release of phosphate. The reaction substrate 2-carboxy-D-ribitol 1,5-bisphosphate (CRBP) was used in place of CA1P as a more easily accessible reagent that is dephosphorylated by CA1Pase at similar rates and has been used as a substitute in other studies (Andralojc *et al*., 2012; Lobo *et al*., 2019). CRBP was added at 3 μL and 20 mM to initiate the reaction with 500 rpm agitation. Reactions were quenched using 40 μL of the colour detection reagent after 10 minutes, unless specified otherwise. One hour after the reaction was quenched, colour development was quantified using a microplate reader for absorption at 640 nm (Supplementary Fig. S3).

### Size exclusion chromatography

An ENrich 650 column (BioRad, #7801650, USA) was equilibrated with 2 column volumes (CVs) of a 50 mM HEPES, pH 8, 200 mM NaCl buffer prior to each run. For runs that included reduced glutathione, 10 mM GSH was included in the buffer by adding 1 mL of 200mM GSH for every 19 mL of buffer. 100-200 μL of purified enzyme (∼20-100 μM) was injected into the column and ran at a flow rate of 0.5 mL min^-1^. Protein Standards (Protein Standard Mix 15 - 600 kDa, #69385, Supelco, USA) were used to quantify protein peak sizes using the same flow rate (Supplementary Table S3). For cases requiring further quantification, diluted fractions were collected at 10 °C in 1.5 mL micro tubes (Sarstedt, Germany, 72.690.001) to quantify protein, TSP, and activity as described above. Some proteins were run at 0.125 mL min^-1^ in attempt to improve peak resolution, however this did not appear to impact peak separation or elution volume. The program findpeaks() from the R package pracma (Borchers 2023) was used to identify all peaks, with visual confirmation (Supplementary Dataset S6).

### Data analyses

Geneious 9.1.8 was used to generate protein alignments for Fig. 1b, Fig. 2a, Fig. 2b, and Fig 5. Further data was processed in R 4.4.2 (R core team, 2024) and RStudio 2025.05.0 (Posit Team, 2025). Data was processed and graphs were prepared using ggplot 2 (Wickham, 2016), agricolae (de Mendiburu, 2023), dplyr (Wickham *et al*., 2023a), readr (Wickham *et al*., 2024), viridis (Garnier *et al*., 2024), pracma (Borchers, 2023), multcomp (Hothorn *et al*., 2008), multcompview (Graves *et al*., 2024), forcats (Wickham, 2023b), FSA (Ogle *et al*., 2025), and rcompanion (Mangiafico, 2025) R packages. For scatter plots, dots represent technical replicates, with mean lines and error bars indicating standard error. All code and data can be found on github (https://github.com/cwnehls/NehlsRamos_etal_JxB_2027).

**Fig. 1.**
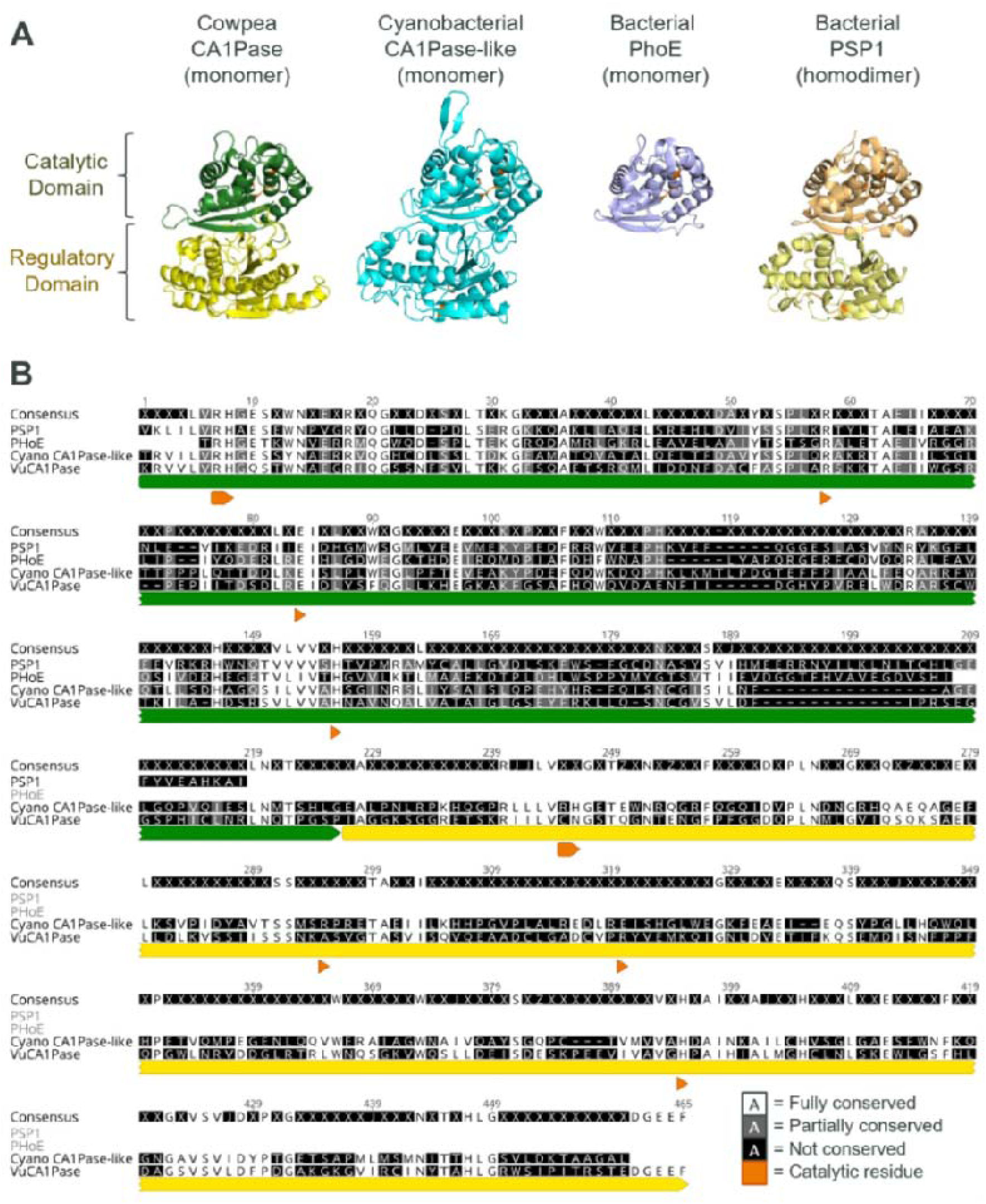
Structural predictions support two distinct domains within CA1Pase. (A) Left- to-right: Predicted AlphaFold 3 structures for cowpea CA1Pase (*Vigna unguiculata*; VuCA1Pase) and a cyanobacterial CA1Pase-like protein from *Myxacorys almenidensis*, crystal structures of a bacterial broad specificity phosphatase (PhoE, 1H2F) from *Geobacillus stearothermophilus*, and a bacterial phosphoserine phosphatase (PSP1, 4IJ5) from *Hydrogenobacter thermophilus*. The predicted transit peptide for VuCA1Pase is removed. Catalytically active residues are highlighted in orange. All structures are shown as monomers, excluding PSP1 which has been resolved as a homodimer. (B) Sequence alignment for the protein sequences of the four enzymes shown, excluding predicted chloroplast transit peptide sequences (Vigun11g130800.1.10, NDJ19632.1, AAK07665.1, BAI68570.1). Sequence alignment and numbering of residues start at the first Lysine (K) of VuCA1Pase. White, grey, and black indicate 100% homology, 50-75%, and not conserved, respectively. The green bar indicates the catalytic domain of VuCA1Pase, yellow bar highlights the regulatory domain. Orange annotations below residue positions denote catalytic residues for phosphatase activity conserved across all four sequences. In the regulatory domain, orange annotations indicate residues that mirror the conserved catalytic residues in the CA1Pase-like sequence.

**Fig. 2.**
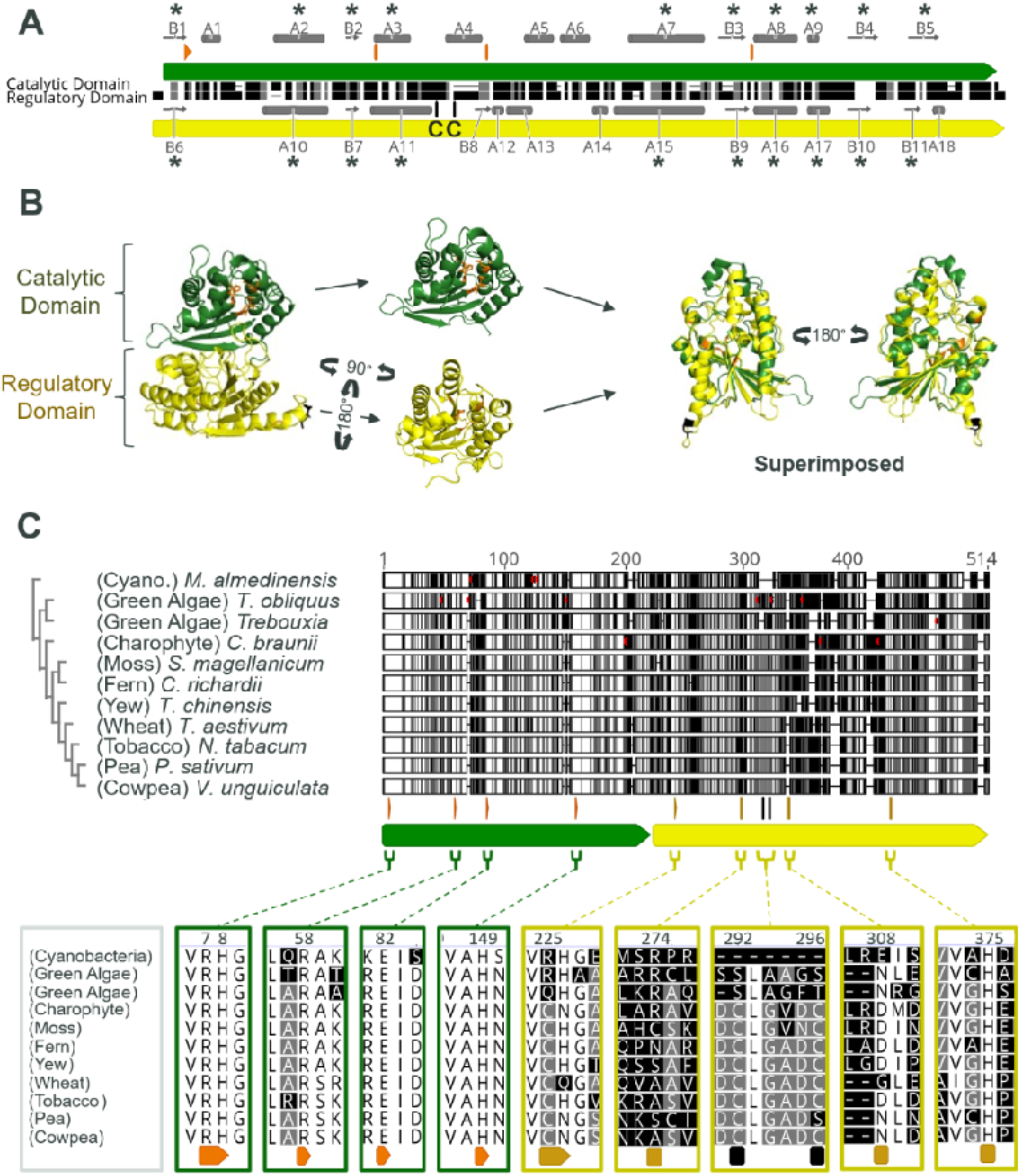
A putative pseudophosphatase regulatory domain mirrors the catalytic domain. (A) Sequence alignment of VuCA1Pase catalytic and regulatory domains. White, grey, and black indicate 100%, 50-75%, and not conserved, respectively. The green bar indicates the catalytic domain of VuCA1Pase, yellow bar highlights the regulatory domain. Black bars above a black “C” indicate the position of the two cysteine residues of the CXXXXC motif. Grey bars indicate secondary structures: arrows indicate beta sheets; cylinders indicate alpha helices. “*” indicate secondary structures conserved in both domains. (B) Comparison of the AlphaFold 3 predicted structures of the VuCA1Pase domains, showing similarity of secondary structures and the position of catalytic residues. Orange indicates catalytic residues and the mirrored residues found in the regulatory domain. (C) Tree alignment of 10 CA1Pase sequences and 1 CA1Pase-like sequence, representing ancestral branches from a wide range of organisms from cyanobacteria to several plant families. Any predicted chloroplast transit peptides were removed, starting alignment and numbering at the lysine of the conserved “KRVV” motif of VuCA1Pase. Green and yellow bars indicate the location of the catalytic and regulatory domains. The positions of catalytic (orange) and mirror (dark yellow) residues, and the putative regulatory cysteine residues (black) are shown in the boxes containing sequence alignment sections at the bottom.

## Results

### CA1Pase domains are structurally similar, with C domain region lacking key phosphatase catalytic residues

To better understand the role and function of the two putative domains of CA1Pase, the structure and sequence of cowpea CA1Pase was compared with related histidine phosphatase family proteins (Fig. 1). These enzymes were screened through structural search by D-I-TASSER (Zheng *et al*., 2026; Supplementary Dataset S4). The two domains previously identified in CA1Pase, a catalytic N domain and C domain (Andralojc *et al*., 2012), resemble those in the predicted cyanobacterial CA1Pase-like structure (Fig. 1a; Supplementary Table S4; Supplementary Table S5). While the catalytic N domain in both proteins strongly resembles the bacterial monomer of PhoE, a broad specificity bacterial phosphatase, their complete monomeric structures parallel the homodimer form of the bacterial phosphoserine phosphatase 1 (PSP1), (Supplementary Table S5). Previously the C domain of CA1Pase was predicted to be a phosphofructokinase domain (Andralojc *et al*., 2012). The NCBI Conserved Domain database, however, identifies the domain as a general histidine phosphatase-like phosphoglycerate mutase (PGM) (COG0406; Lu *et al*., 2020).

For all four proteins, the known catalytic residues - two arginines (R), two histidines (H), and one glutamate (E) - are conserved in the catalytic domains (Fig. 1B). These are conserved in the C domain for the cyanobacterial CA1Pase-like enzyme, but not in the C domain for VuCA1Pase, which retains only the last histidine, H375. While similar to the structure of the catalytic domain of PSP1, the C domain has several differences: an extended alpha helix on the side facing the catalytic domain opening and a lack of conserved alpha helices in the region mirroring the catalytic pocket of the cyanobacterial C domain (Fig. 1A). The structural similarity that is retained may allow for allosteric binding of ligands in the predicted pseudophosphatase pocket (Supplementary Fig. S4). Due to this and the expected lack of catalytic capacity, we investigated if the C domain may function as a regulatory domain.

### Conserved domain structures suggest an ancient gene duplication and fusion event

To identify key and unique regions of the regulatory domain, the two domains were investigated via superimposition of predicted structures and AlphaFold sequence alignments. To then establish key residues and potential ancestral motif shifts within these domains, a selection of CA1Pase and CA1Pase-like sequences were identified by BLAST from organisms ranging from cyanobacteria to plants for comparison.

Initial alignment of the two domains by sequence and secondary structure reinforced observed similarities of the regulatory domain and the C domain of the cyanobacterial sequence (Fig. 2A). While the pairwise identity of residues between the domains is low (∼23%), greater identity is conserved in the secondary structure. All five beta sheets in the catalytic domain are conserved in the regulatory domain along with five of nine alpha helices. Superimposing the structures reveals high conservation in the position of these secondary structures (Fig. 2B). Interestingly, shortly after the extended alpha helix (A11) is a conserved cysteine pair motif (CXXXXC) predicted to form a disulphide bond (Fig. 1A-B). This motif is not present in the catalytic domain, suggesting a unique potential redox role for the regulatory domain.

A phylogenetic tree comparing the CA1Pases identified by BLAST highlighted high conservation of the catalytic domain active site residues from cyanobacteria through to land plant lineages (Fig. 2C). This was not true for the equivalent residues in the regulatory domain: the first arginine is only conserved in green algae; the first catalytic histidine is retained only in certain lineages down to flowering plants; the second arginine is lost with the onset of land plants; the key glutamate is present solely in the cyanobacterial sequence. Only the previously mentioned second histidine, H375, is conserved throughout all lineages. The CXXXXC motif is highly conserved beginning with the charophytic, suggesting redox regulation for CA1Pase may originate within Streptophytes. The lack of conserved residues across the regulatory domain, combined with the less well predicted structure of the region that mimics the catalytic substrate binding pocket, may suggest that substrate binding specificity could be tied to individual lineages. This may, in turn, allosterically regulate CA1Pase activity depending on species or lineage differences in stromal metabolite content.

### Structural predictions and *in vitro* analyses suggest the regulatory domain influences CA1Pase dimer formation

Several enzymes in the chloroplast form multimeric structures that can regulate catalytic activity (Podesta *et al*., 1990; Yu *et al*., 2020; Waheeda *et al*., 2023). To investigate potential regulatory oligomer formation, predicted multimeric structures for mature VuCA1Pase were generated and stable forms identified (Supplementary Fig. S1). Beyond the monomer, the homodimer was predicted to be the most stable form. Sequences for CA1Pases from 252 species were aligned to identify well conserved residues that may support dimer formation (Supplementary Dataset S1). Residues conserved in over 95% of those sequences are highlighted in the predicted dimer structure (Fig. 3A). Of the conserved residues in the regulatory domain, the majority are located in the predicted dimer interface, suggesting a well conserved ability to form dimers across flowering plants i*n vivo*. The previously identified CXXXXC motif was also found to be part of this dimer interface (Fig. 3B). The predicted surface facing position of the disulphide bond suggests a redox role in dimer formation and stabilisation.

**Fig. 3.**
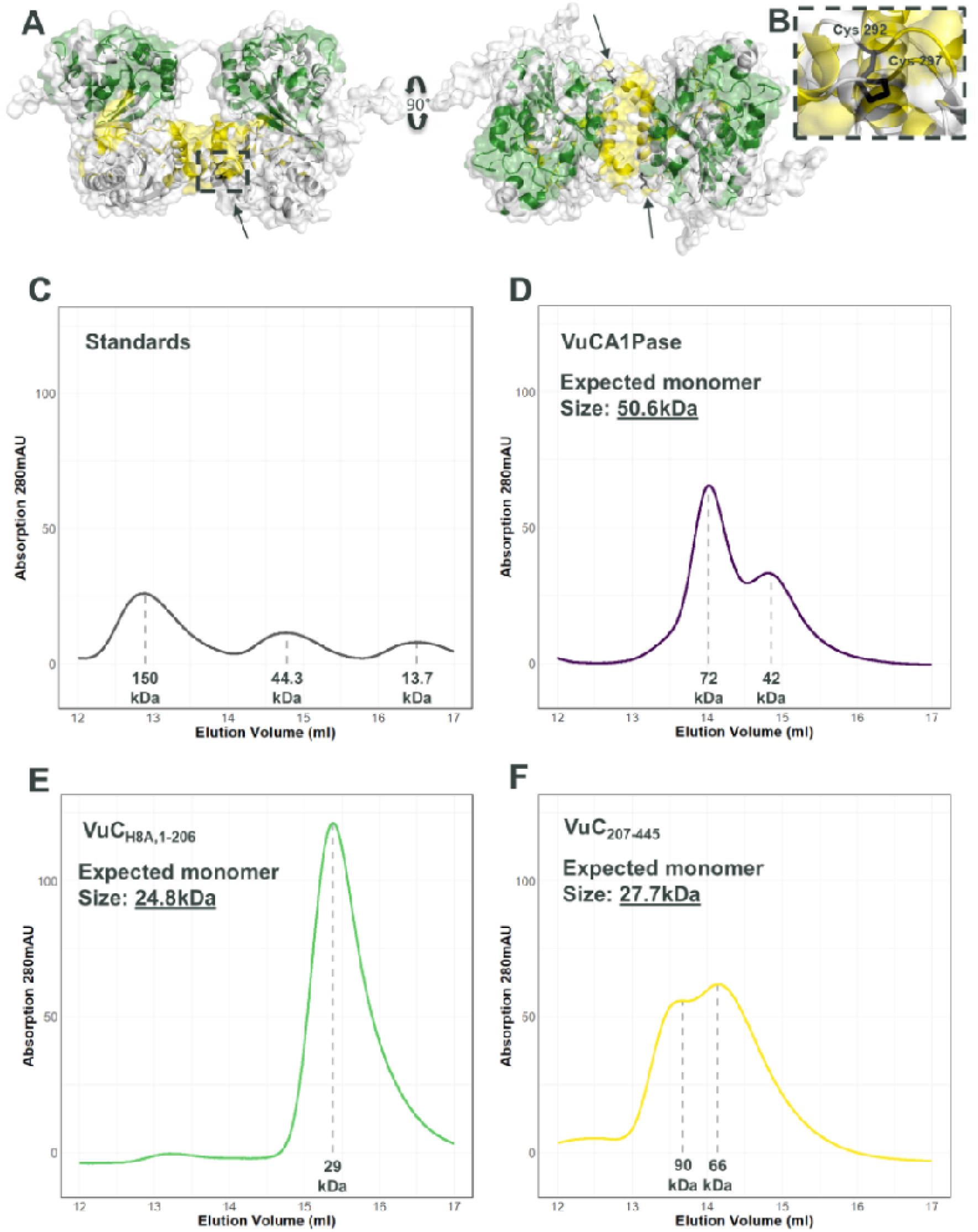
The regulatory domain of CA1Pase is required for dimer formation. (A) Side and top view of the AlphaFold 3 predicted dimer structure of VuCA1Pase. Coloured regions represent residues conserved in >95% of 252 CA1Pase sequences, ranging from monocots to dicots. Green and yellow represent conserved catalytic and regulatory domain residues, respectively. The presence of the predicted disulfide bond formation is shown within the black box and indicated by black arrows. (B) A zoom into the redox motif region shown in the dotted line box in Figure 3A. Cysteines 292 and 297 are shown along with the presence of the predicted disulfide bond. This motif neighbours conserved residues that are part of the predicted dimer interface. (C-F) Size exclusion chromatography (SEC) traces for (C) protein standards (γ-globulins, Ovalbumin, Ribonuclease A type I-A), (D) VuCA1Pase, (E) VuCA1Pase lacking the regulatory domain (VuC_H8A,1-206_), and (F) VuCA1Pase lacking the catalytic domain (VuC_207-445_). Estimated size in kilodaltons (kDa) is indicated below each peak, shown by a dotted grey line.

When analysed by size exclusion chromatography, recombinant VuCA1Pase repeatedly showed two peaks with sizes consistent with a monomer (near 40kDa) and a larger peak that may be a dimer (near 70kDa; Fig. 3D). While the peaks are estimated to be smaller than the near 50 and 100kDa expected, sizes determined by size exclusion are not exact and can be influenced by a variety of factors including the shape of the protein (Fekete *et al*., 2014; Burgess, 2018). To assess the ability of each domain to independently form multimers, the domains were expressed separately as truncated proteins. The catalytic domain variant VuC_H8A,1-206_, showed consistent peaks near their monomer size (Fig. 3E), while the regulatory domain variant, VuC_207-445_ showed peaks that were consistent with multimeric sizes, typically approximating trimers and dimers (Fig. 3F). Given the regulatory domain uniquely shows affinity for oligomer formation, this further supports a role in dimer formation for VuCA1Pase *in vivo*.

### Two key regulatory domain cysteines influence redox sensitive CA1Pase activity and dimer formation

The combined roles of dimer formation and redox regulation of the regulatory domain were investigated through the generation of recombinantly expressed cysteine mutants, whereby one or both key cysteines (C292, C297) were replaced with alanine (Supplementary Fig. S5). CA1Pase activity of these mutants was then compared to the recombinant wild type VuCA1Pase across a gradient of reducing conditions. From 100% oxidised glutathione (GSSG) to 100% reduced glutathione (GSH), unmodified VuCA1Pase showed a distinct increase in activity in reducing conditions (Fig. 4A). This supports previous findings in tobacco and French bean which showed a stimulatory effect of GSH, and inhibitory effect of GSSG (Holbrook *et al*., 1991; Heo & Holbrook, 1999). The reducing conditions also appeared to shift the oligomeric state of VuCA1Pase, favouring the dimer form in the presence of 5mM DTT (Fig. 4A) and GSH (Supplementary Fig. S6). All cysteine mutants showed a lower activity in fully reducing conditions compared to the wild type (Fig. 4B-D), with those containing the C297A mutation showing an inhibitory effect when fully reduced (Fig. 4C-D). The C292A mutation alone showed an unexpected increase in activity from oxidised to reducing conditions despite the presumed loss of disulphide formation potential (Fig. 4B).

**Fig. 4.**
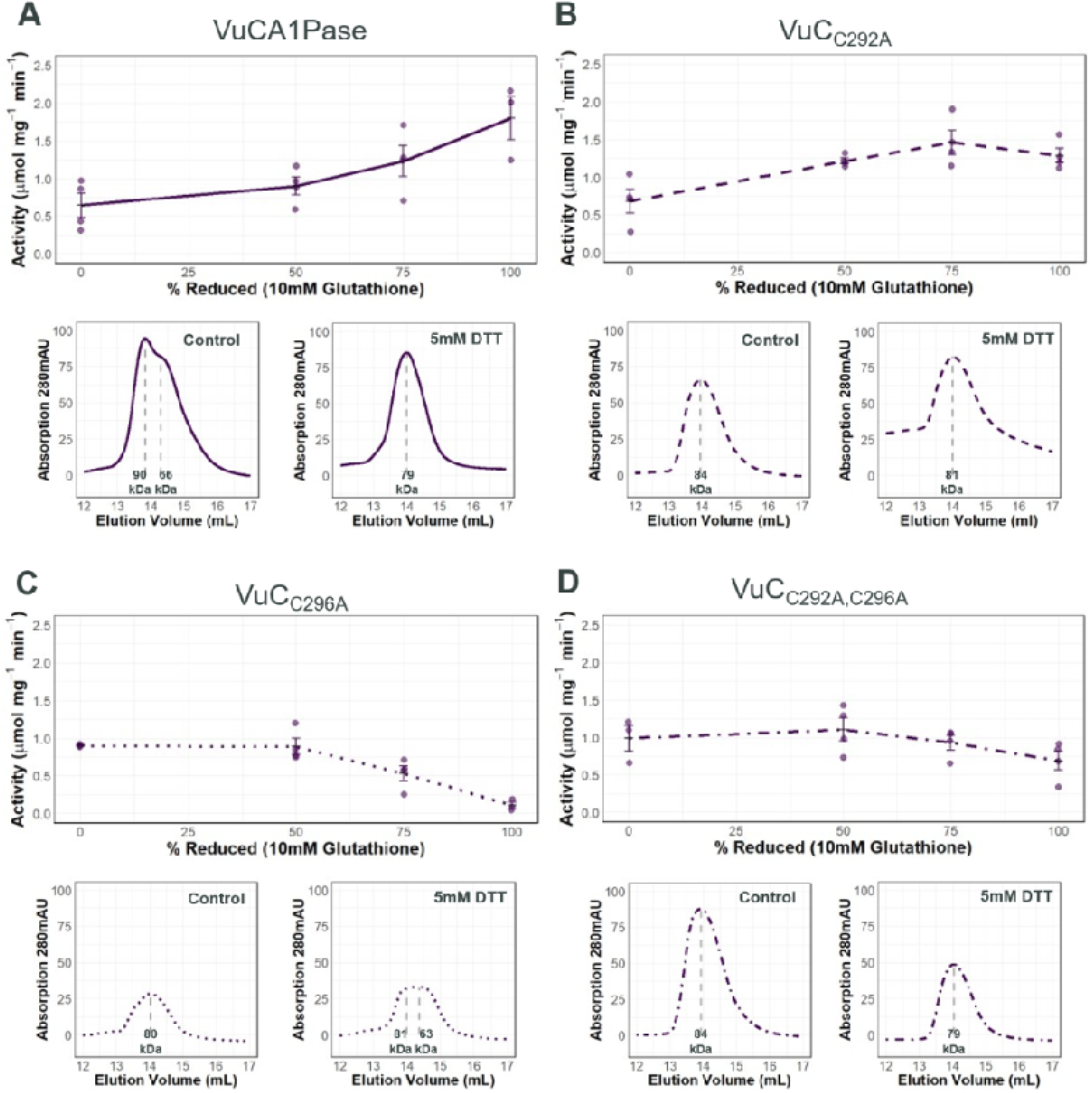
Redox regulation of CA1Pase is linked to dimer formation. (A-D) Graphs at the top of each panel display CA1Pase specific activity in presence of increasing concentrations of reduced glutathione (GSH, 0-100%). Graphs below show the traces of size exclusion chromatography in the absence (left) or presence of 5 mM DTT reducing agent (right), with predicted peak sizes. (A) VuCA1Pase, (B) single cysteine mutant VuC_C292A_, (C) single cysteine mutant VuC_C297A_, (D) double cysteine mutant VuC_C292A,C297A_. Technical replicates for each GSH treatment (n) = 3-4. Identified peaks are denoted by a grey dotted line underneath.

For all the cysteine mutants (Fig. 4B-D), no monomer peak was seen in non-reducing conditions. With VuC_C297A_, reducing conditions resulted in a near total loss of activity along with the presence of a monomer sized peak. This monomer peak could relate to the loss of activity. Results from *in vitro* activity of the individual peaks of wild type VuCA1Pase supports the dimer as the more active variant (Supplementary Fig. S7). Why the C297A variant would so drastically lose activity in highly reducing conditions is unclear.

### Variation in two key cysteines is correlated with low Rubisco dark inhibition in Fabaceae

To identify the potential impact of the CA1Pase redox regulation, natural variants of CA1Pase were identified lacking one or both key cysteines in species where dark inhibition data is available (Supplementary Table S2; Nehls-Ramos *et al*., 2026). While cysteine mutants were found across angiosperms, a more distinct trend was seen in Fabaceae, the most well sampled family (Fig. 5; Nehls-Ramos *et al*., 2026). For the bean family, fifteen species were identified that had both data for dark inhibition and a known CA1Pase sequence. Of these, five had serine or threonine in place of one or both of C292 and C297 (Fig. 5). All five had low accumulation of dark inhibitors despite other related species having much higher accumulation. This trend suggests CA1Pases lacking redox regulation capacity from the CXXXXC motif may retain activity regardless of light or stress related redox state, mirroring our *in vitro* cysteine mutant work. It is therefore possible that these CA1Pase variants may continue to degrade inhibitors such as CA1P during night and/or dark conditions.

**Fig. 5.**
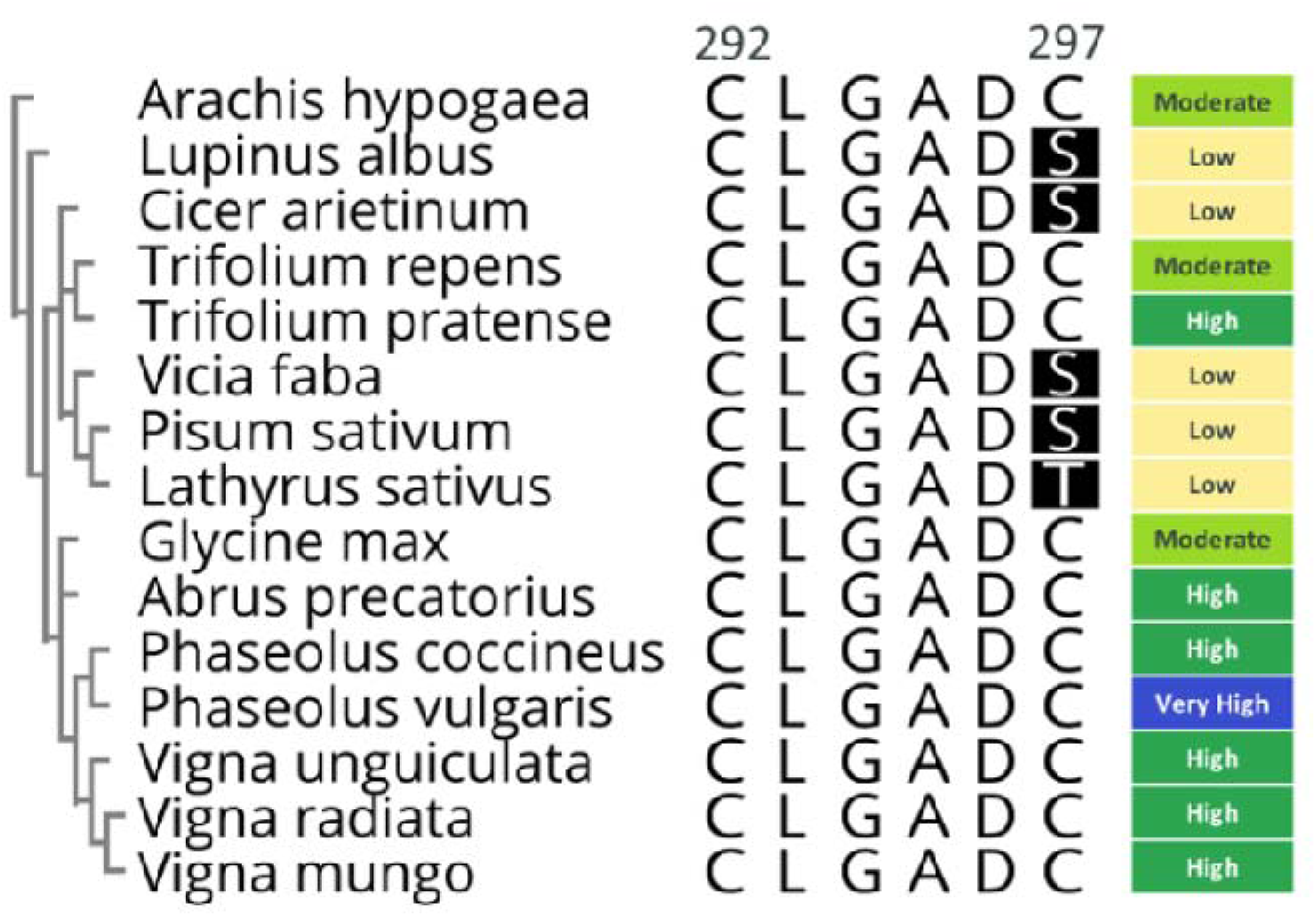
C297 variants in Fabaceae CA1Pases correlate with low Rubisco dark inhibition. Alignment of mature CA1Pase from 15 Fabaceae species, ordered by tree alignment (left), highlighting conservation of the CXXXXC motif. Species represent those from Fabaceae with both sequence and dark inhibition data available (see Nehls-Ramos *et al*., 2026). Positions of the regulatory domain CXXXXC motif (C292 & C297) identified in *Vigna unguiculata* are annotated. Residues are numbered as described in Materials and Methods. Black highlight indicates residue differences from consensus. Coloured boxes at right indicate known prevalence of dark inhibition, as classified by Nehls-Ramos *et al*. (2026).

## Discussion

CA1Pase plays an important role in dephosphorylating certain sugar phosphate derivatives that inhibit the activity of Rubisco. We used a combination of *in silico* and *in vitro* protein characterisation to elucidate the function and mechanisms of CA1Pase regulation in the legume crop cowpea. Our results suggest an ancient monomeric origin for this enzyme, later developing capacity for dimerisation and conversion of the C domain into a pseudophosphatase regulatory domain in Viridiplantae. Appearance of a key CXXXXC motif suggests redox regulation arose in the Charophytes. We found evidence of CA1Pase forming redox regulated homodimers, which may confer heightened activity compared to the monomer form. This redox regulation was eliminated *in vitro* through mutating one or both cysteines, and our results suggest an association between natural cysteine variants of CA1Pase and previously identified low dark inhibitor accumulation in some plant species.

While previously the evolutionary origins of CA1Pase have been difficult to parse, comparison with its bacterial and cyanobacterial relatives enables a clearer picture. In histidine phosphate phosphoglycerate mutase (PGM) family enzymes, a single phosphatase domain is sufficient, as with PhoE enzyme (Fig. 1A). In some cases, these enzymes form dimers or higher order homooligomers as with PSP1, which can improve activity and stability of the enzyme complex (Fig. 1A; Marianaya *et al*., 2004). It is therefore not so surprising that this type of phosphatase has the potential to form a fused dimer via a gene fusion duplication event that could retain this stable form (Yanai *et al*., 2002; Muthahari *et al*., 2025). Such events are not uncommon and can provide diversification of domain roles, while also improving the stability of a potentially higher activity structure (Rullán-Lind *et al*., 2018). It is possible that the binding of ligands to one of the catalytically active domains may also help to regulate the activity of the opposite domain allosterically, providing a possible sensor function to further regulate activity of this phosphatase-like enzyme. For example, such allosteric regulation occurs with phosphofructokinase (Compton & Patrick, 2025). This fusion event in CA1Pase likely occurred in cyanobacteria prior to the endosymbiotic origin of chloroplasts or was transferred to an early chloroplast via horizontal gene transfer. This is supported by the ubiquity of CA1Pase-like sequences across cyanobacterial classes (Supplementary Dataset S1).

The redox-sensitive dimerisation function of the regulatory domain in land plants may have first occurred in charophytes, with the development of the extended alpha helix 11, and the nearby CXXXXC motif (C292, C297) that is absent in chlorophytic and cyanobacterial CA1Pase-like sequences (Fig. 2A; Fig. 2C; Supplementary Dataset S1). Despite this, the predicted structures for the chlorophyte algae *Tetradesmus obliquus* and *Trebouxia* suggest dimerisation occurring in this division, and that this potentially arose prior to gaining the redox sensitive trait (Supplementary Dataset S1; Supplementary Dataset S2). Regulation of day-night dephosphorylation of specific metabolites in early land plants and their charophytic ancestors may have driven an adaptation of CA1Pase.

Given the loss of catalytic residues in the regulatory domain appears to have occurred in both chlorophytes and charophytes, and that the regulatory domain pocket faces towards the proposed dimer interface, the regulatory domain may have first functioned as a pseudophosphatase sensor for certain sugar-phosphate metabolites in the chloroplast, potentially linked to dimer interactions (Fig. 3A; Reiterer *et al*., 2020). This function is supported by the evidence of various sugar phosphate CBB metabolites, such as FBP, enhancing CA1Pase activity *in vitro* in previous studies, along with *in silico* predictions suggesting certain chloroplast metabolites may interact with the regulatory pocket (Table 1; Supplementary Fig. S4). These predicted structures however do not give the complete picture and are limited to the related resolved structures they are based on. The resolved structure of land plant CA1Pase with FBP would help elucidate this potential interaction, and any impacts it may have on the proposed dimer structure.

In agreement with previous *in vitro* studies, cowpea CA1Pase also appears to be redox sensitive, with reducing conditions enhancing activity (Table 1; Fig. 4A). Here we show that the disruption of one or both key cysteines in the regulatory domain, C292 and C297, either decreases or reverses the positive effect on CA1Pase activity. This is likely tied to the disruption of the formation of the more active dimer form (Fig. 3B-D). The cysteine redox state impact on dimer formation may be due to this covalent disulphide bond providing a more rigid loop structure, hindering more stable conformations for the dimer interface (Fig. 3A). Currently no experimental CA1Pase structures are available; future structures of the reduced and oxidised forms would help to better elucidate this mechanism.

Between the two cysteines, C297 appears the more important residue for disrupting dimer formation. Mutating this residue to alanine resulted in a distinct loss of activity in reducing conditions that was not seen with the C292A variant (Fig. 4B-D). This may be due to C297 being more solvent available, allowing for further interactions by this residue when forming disulphide bonds with the dimer interface that is clearly seen in the predicted structure. Reduced glutathione can form mixed disulphides with cysteine by S-glutathionylation, which may explain the partial increase of activity with the C292A variant that retained C297 (Heo & Holbrook, 1999; Musaogullari & Chai, 2020). Formation of mixed disulphides for C297 may result in a conformation that further stabilizes dimer formation, given it does not directly interact with the dimer interface to cause a disruption by disulphide formation. The opposite may be true for C292 which may disrupt the dimer interface when converted to a mixed disulphide. An explanation for the loss of activity in the dual mutant C292A,C297A in reducing conditions is less clear, potentially suggesting the slight deleterious effect of mixed disulphides in other parts of the CA1Pase structure for solvent available cysteines. A more active CA1Pase may be possible by retaining the hydrophobic C292 while exchanging C297 for a hydrophilic residue. This could result in a conformation of this region that further stabilises the dimer interface.

Across plant species, some CA1Pase cysteine variants of C297 possess hydrophilic, polar uncharged residues at this position, such as serine or threonine (Fig. 5). These amino acids are encoded by codons with a single nucleotide different from cysteine, requiring only a single nucleotide mutation to occur. These species are known to have low Rubisco dark inhibition, potentially indicating increased inhibitor degradation in dark conditions due to lacking a low-activity oxidised state. This trend of sudden phylogenetic loss of dark inhibition had been previously identified in our work and can now be consistently linked with these cysteine variants within the Fabaceae (Nehls-Ramos *et al*., 2026). Similar evidence points to this link in the limited data available for Brassicaceae. It should be noted that there is limited current information on the expression and abundance of CA1Pase in these species, which would further elucidate this lower dark inhibition across species.

Given crops regularly face stress conditions which in some species has been shown to increase production of Rubisco inhibitors (Parry *et al*., 2008; Carmo-Silva *et al*., 2010), the downregulation of CA1Pase activity in periods of decreased photosynthesis and elevated stromal ROS means a decreased potential for Rubisco inhibitor degradation (Slattery *et al*., 2018). This could be relevant during heat and drought stress conditions, where both ROS and Rubisco inhibitor generation is expected to be higher (Parry *et al*., 2008; Carmo-Silva *et al*., 2010; Slattery *et al*., 2019). Such inhibitors include those produced as misfire products from increased oxygenation rates (Pearce, 2006; Parry *et al*., 2008; Orr *et al*., 2023). A CA1Pase that is resistant to the impacts of oxidising dark and stress conditions could mean continued inhibitor degradation regardless of environment. Increased CA1Pase activity may also help with the recycling of phosphate from other inhibitory compounds such as PDBP, supporting sustained chloroplast phosphate homeostasis. Regular degradation of inhibitors may be essential for sustaining available phosphate for the light reactions of photosynthesis (Leister *et al*., 2023). The direct benefits of the proposed redox insensitive CA1Pase cysteine variants in some Fabaceae remains to be explored and may further elucidate the utility and role of this trait (Fig. 5).

CA1Pase contributes to regulating the abundance of inhibitors and the activity of Rubisco. Our investigation of the structure and function of cowpea CA1Pase, identified evolutionary trends and highlighted a key CXXXXC motif that impacts redox regulation. The physiological impact of altering the abundance of such inhibitors through manipulating the abundance and activity of CA1Pase remain to be fully elucidated *in planta*. Future studies are warranted to interrogate and verify the specific mechanism for sugar phosphate allosteric binding in the regulatory domain and redox-dependant dimerisation via experimentally resolved structures. Metabolic, proteomic, and physiological studies, combined with modified expression of CA1Pase in species presenting low and high Rubisco dark inhibition may further elucidate the combined role of CA1P and CA1Pase as regulators of carbon assimilation by Rubisco as well as any ancillary role in the metabolic regulation of the chloroplast. Through identifying new avenues for improved Rubisco inhibitor degradation, we can gain further insight into optimisation for improved photosynthetic activity towards maximising crop resilience and output.

## Supplementary data

**Supplementary Fig. S1.** AlphaFold 3 predicted structures with confidence indicated

**Supplementary Fig. S2.** SDS-PAGE gels of purified VuCA1Pase and variants

**Supplementary Fig. S3.** Phosphate detection diagram

**Supplementary Fig. S4.** Alphafold 3 predicted structures for regulatory domain ligand binding

**Supplementary Fig. S5.** Labelled AlphaFold 3 predicted structures for VuCA1Pase cysteine mutants

**Supplementary Fig. S6.** Effect of glutathione on SEC elution peaks for VuCA1Pase and VuC_C292A,C297A_

**Supplementary Fig. S7.** VuCA1Pase eluted fraction activity and SDS gel bands

**Supplementary Table S1.** Full list of species with known CA1Pase sequences and dark inhibition levels

**Supplementary Table S2.** Recombinant protein purity and date of purification

**Supplementary Table S3.** Size exclusion column protein elution expected protein sizes with treatments

**Supplementary Table S4.** Pairwise identity of aligned peptide sequences

**Supplementary Table S5.** Root mean square deviation of aligned predicted and resolved structures

**Supplementary Dataset S1.** Geneious alignments and sequences

**Supplementary Dataset S2.** AlphaFold 3 predicted structures

**Supplementary Dataset S3.** Crystal structures for broad specifcity phosphatases

**Supplementary Dataset S4.** D-I-Tasser predicted protein structures

**Supplementary Dataset S5.** *in vitro* data for biochemical assays

**Supplementary Dataset S6.** SEC runs for CA1Pases and variants

## Acknowledgements

The authors thank colleagues in the Photosynthesis team at Lancaster for useful discussions and data interpretation, especially Dr. Armida Gjindali, Dr. Joana Amaral, Dr. Clayton Dilks, Dr. Felix Townsend, and Dr. Dawn Worrall. We also thank Dr. Mitchell Altschuler (University of Illinois at Urbana-Champaign) for useful discussions.

## Author contributions

CN, ECS, and DJO: conceptualisation; CN: data curation; CN: formal analysis; ECS and DJO: funding acquisition; CN: investigation; CN, ECS and DJO: methodology; ECS and DJO: resources; ECS and DJO: supervision; CN: visualisation; CN: writing - original draft; CN, ECS, and DJO: writing - review & editing.

## Conflict of interest

None declared.

## Funding

This work was supported by the project Realizing Increased Photosynthetic Efficiency (RIPE), that is funded by Gates Agricultural Innovations grant investment 57248, awarded to Lancaster University by the University of Illinois, USA. CNR is also supported by a Lancaster Environment Centre (LEC) Postgraduate Research Studentship from Lancaster University.

## Data availability

The data collated and generated for this paper is included in the excel file found in (https://github.com/cwnehls/NehlsRamos_etal_JxB_2027). Within the github file the code used for analyses and figure generation are included.

## Abbreviations

CA: 2-carboxy-D-arabinitol
CA1P: 2-carboxy-D-arabinitol 1-phosphate
CA1Pase: 2-carboxy-D-arabinitol 1-phosphate phosphatase
CBB: Calvin-Benson-Basham
CRBP: 2-carboxy-D-ribitol-1,5-bisphosphate
cTP: chloroplast transit peptide
FBP: fructose 1,6-bisphosphate
GSH: reduced glutathione
GSSG: oxidised glutathione
PDBP: D-glycero-2,3-pentodiulose-1,5-bisphosphate
PGM: phosphoglycerate mutase
PSP1: phosphoserine phosphatase 1
Rca: Rubisco activase
ROS: Reactive oxygen species
RuBP: ribulose 1,5-bisphosphate
TSP: total soluble protein
VuCA1Pase: *Vigna unguiculata* 2-carboxy-D-arabinitol 1-phosphate phosphatase

## Notes

### Competing Interest Statement

The authors have declared no competing interest.

